# Molecular Detection of SARS-CoV-2 in Formalin Fixed Paraffin Embedded Specimens

**DOI:** 10.1101/2020.04.21.042911

**Authors:** Jun Liu, April M. Babka, Brian J. Kearney, Sheli R. Radoshitzky, Jens H. Kuhn, Xiankun Zeng

**Author notes:** Correspondence (X.Z.): United States Army Medical Research Institute of Infectious Diseases (USAMRIID), 1425 Porter Street, Fort Detrick, Frederick, MD 21702, USA.

## Abstract

Severe acute respiratory syndrome coronavirus 2 (SARS-CoV-2), the cause of human coronavirus disease 2019 (COVID-19), emerged in Wuhan, China in December 2019. The virus rapidly spread globally, resulting in a public-health crisis including more than one million cases and tens of thousands of deaths. Here, we describe the identification and evaluation of commercially available reagents and assays for the molecular detection of SARS-CoV-2 in infected formalin fixed paraffin embedded (FFPE) cell pellets. We identified a suitable rabbit polyclonal anti-SARS-CoV spike protein antibody and a mouse monoclonal anti-SARS-CoV nucleocapsid protein (NP) antibody for cross detection of the respective SARS-CoV-2 proteins by immunohistochemistry (IHC) and immunofluorescence assay (IFA). Next, we established RNAscope *in situ* hybridization (ISH) to detect SARS-CoV-2 RNA. Furthermore, we established a multiplex fluorescence ISH (mFISH) to detect positive-sense SARS-CoV-2 RNA and negative-sense SARS-CoV-2 RNA (a replicative intermediate indicating viral replication). Finally, we developed a dual staining assay using IHC and ISH to detect SARS-CoV-2 antigen and RNA in the same FFPE section. These reagents and assays will accelerate COVID-19 pathogenesis studies in humans and in COVID-19 animal models.

## Introduction

Severe acute respiratory syndrome coronavirus 2 (SARS-CoV-2), the etiologic agents of human coronavirus disease 2019 (COVID-19), initially emerged in Wuhan, Hubei Province, China in December 2019 (1-3). As of April 9, 2020, 1,436,198 cases of COVID-19, including 85,522 deaths have been reported worldwide (4).

SARS-CoV-2 has a nonsegmented, linear, positive-sense, multicistronic genome and produces enveloped virions (5). The virus is classified as a betacoronavirus (*Nidovirales*: *Coronaviridae*) together with the other two highly virulent human pathogens severe acute respiratory syndrome coronavirus (SARS-CoV) and Middle East respiratory syndrome coronavirus (MERS-CoV) (6). The SARS-CoV-2 genomes shares 79.6% and 50.0% nucleotide sequence identity with the genomes of SARS-CoV and MERS-CoV, respectively (5). Similar to SARS-CoV, SARS-CoV-2 virions use their spike (S) glycoproteins to engage host-cell angiotensin I-converting enzyme 2 (ACE2) to gain entry into host cells and host-cell transmembrane serine protease 2 (TMPRSS2) for S priming (7).

Bats are speculated to be the natural reservoir of SARS-CoV-2 because numerous other betacoronaviruses are of chiropteran origin (8, 9). However, although the COVID-19 pandemic may have begun with a bat-to-human transmission event, it appears that close to all human infections trace back to respiratory droplets produced by infected people and fomites (respiratory droplet landing sites) (10, 11). Human infections lead to various degrees of disease severity, ranging from asymptomatic infection or mild symptoms to fatal pneumonia. Older patients or patients with chronic medical conditions are more vulnerable to becoming critically ill with poor prognosis (12). The most common symptoms and clinical signs of COVID-19 are fever, cough, dyspnea, and myalgia with medium incubation period of 4 days (13-15). Ground-glass opacity is the most common radiologic finding on chest CT upon admission (13-15). Bilateral diffuse alveolar damage, alveolar hemorrhage and edema, interstitial fibrosis and inflammation, and type II pneumocyte hyperplasia are observed in post-mortem human lungs (16-18).

At the time of writing, there are no animal models that truly mimic the disease spectrum and pathogenesis of COVID-19. However, small animals (e.g., human ACE2 transgenic laboratory mice (19), cats (20), domestic ferrets (20, 21), golden hamsters (22)), and nonhuman primates (e.g., rhesus monkeys (23, 24), crab-eating macaques (25)), are used to study SARS-CoV-2 infection as alveolar damage, interstitial inflammation, and viral shedding occur in these animal models to various degree. It is hoped that further development of these and other animal models will help overcome the current roadblock to evaluating the efficacy of candidate medical countermeasures (MCMs) against and the pathogenesis of COVID-19.

Detection of viral antigen using IHC or IFA techniques and detection of viral nucleic acids using ISH within infected, but inactivated, human or animal model tissues greatly facilitates detection of viral infection and thereby pathogenesis and MCM efficacy studies. These techniques become paramount in particular for studies of a potential pathogen that does not cause overt, or causes only mild, disease, such as SARS-CoV-2 in the currently available animal models. Viral antigen-based immunostaining has been used to detect SARS-CoV-2 antigen in both post-mortem human and animal tissues (1, 16, 22, 25). However, the antibodies used in these studies were produced in-house and therefore are not commonly available. Identification and characterization of commercially available anti-SARS-CoV-2 antibodies and ISH assays that can be used to detect SARS-CoV-2 in FFPE tissues are therefore critically needed.

Here, we describe the evaluation of a rabbit polyclonal anti-SARS-CoV S antibody and a mouse monoclonal anti-SARS-CoV NP antibody that are commercially available and, in IHC and IFA, recognized respective SARS-CoV-2 proteins in FFPE specimens. We also identify two commercially available ISH assays that can be used to efficiently detect SARS-CoV-2 RNA in such specimens and develop a dual staining assay using IHC and ISH to detect SARS-CoV-2 S and RNA in the same FFPE section.

## Results

### Identification of antibodies suitable for detection of SARS-CoV-2 by IHC and IFA in FFPE specimens

To identify antibodies that can be used to detect SARS-CoV-2 in human and animal tissues, we searched for commercially available SARS-CoV antibodies that recognize epitopes that are likely conserved in SARS-CoV-2. We identified six antibodies, including three rabbit polyclonal antibodies, against SARS-CoV S, one rabbit polyclonal antibody against SARS-CoV nucleocapsid protein (NP), and one rabbit and one mouse monoclonal antibody against SARS-CoV NP that may cross react with SARS-CoV-2 (Supplemental Table 1). Additionally, we also identified a rabbit monoclonal antibody against SARS-CoV-2 S (Supplemental Table 1). To evaluate whether these six antibodies can recognize SARS-CoV-2 in FFPE specimens, we performed IHC on FFPE pellets of Vero 76 cells infected with SARS-CoV-2. We identified one rabbit polyclonal antibody against SARS-CoV S (Sino Biological, Chesterbrook, PA, USA; #40150-T62-COV2) and a mouse monoclonal antibody against SARS-CoV NP (Sino Biological, 40143-MM05) that did not stain uninfected, but stained SARS-CoV-2-infected FFPE cell pellets (Figure 1A–D). Furthermore, we did IFA using these two antibodies. Interestingly, in comparison to relatively concentrated detection of SARS-CoV-2 NP (red) in cytoplasmic membrane, the SARS-CoV-2 S (green) is more confined in perinuclear inclusion bodies (Figure 1E).

**Figure 1.**
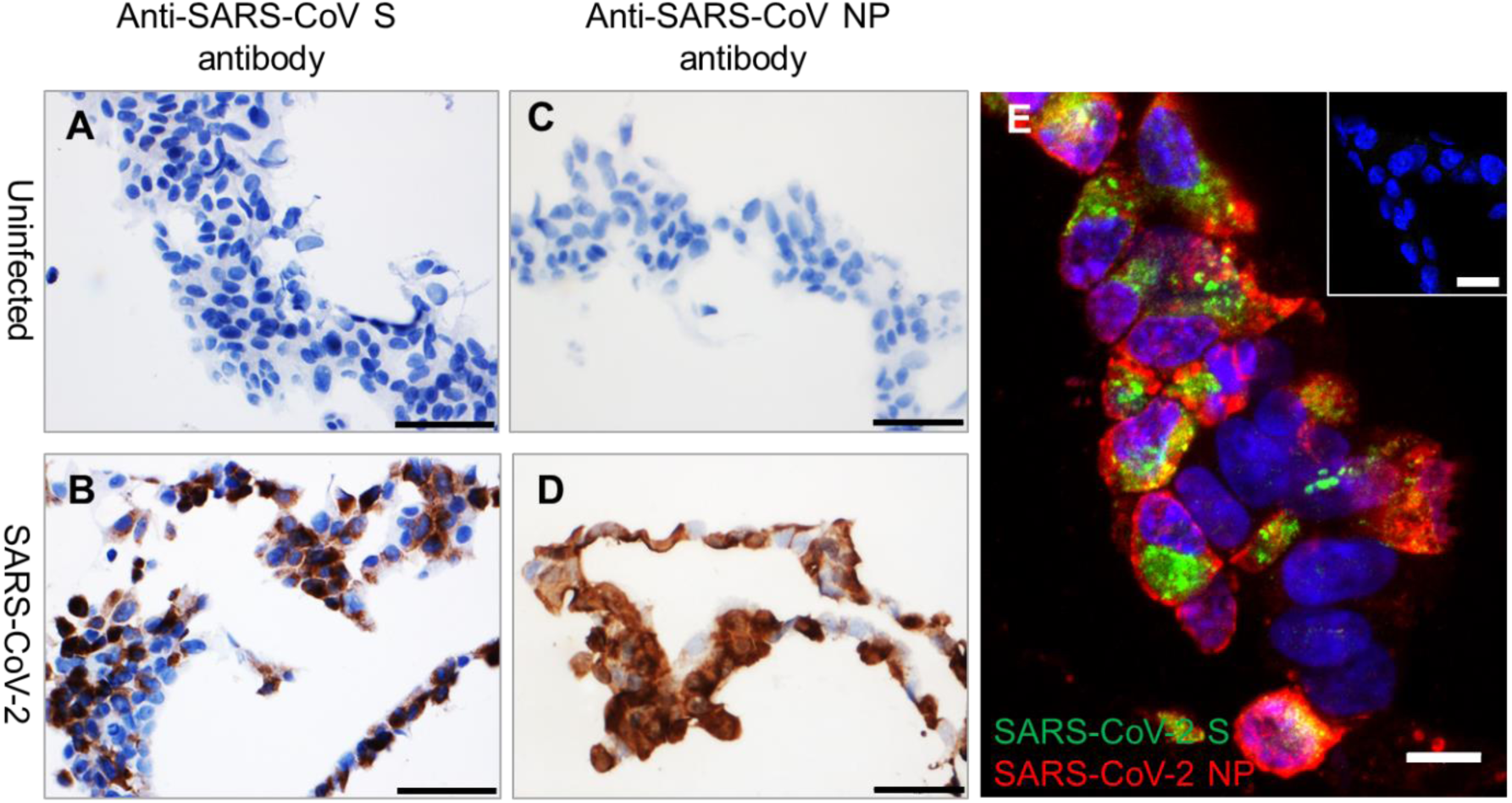
Detection of SARS-CoV-2 antigens by IHC and IFA in FFPE cell pellets. (**A–B**) In comparison to uninfected control FFPE cell pellets (**A** and **C**), SARS-CoV-2 S (brown, **B**) and SARS-CoV-2 NP (brown, **D**) can be detected in FFPE SARS-CoV-2-infected cell pellets. Nuclei are stained blue (hematoxylin). (**E**) Immunofluorescence staining to detect SARS-CoV-2 S (green) and NP (red) in FFPE SARS-CoV-2-infected cell pellets. Inset of (**E**) is uninfected control FFPE cell pellets. Nuclei are stained blue (DAPI). Scale bar, 50 μm in (**A**–**D**), 20 μm in inset of (**E)**, and 10 μm in (**E)**.

### Detection of SARS-CoV-2 RNA by ISH in FFPE tissues

We have previously reported the development of RNAscope ISH assays to detect various high-consequence viruses including Ebola virus (EBOV; *Filoviridae*: *Ebolavirus*), Marburg virus (MARV; *Filoviridae*: *Marburgvirus*), Lassa virus (LASV; *Arenaviridae*: *Mammarenavirus*), and Nipah virus (NiV; *Paramyxoviridae*: *Henipavirus*) in FFPE animal tissues (26-29). Here we report the successful use of the RNAscope ISH assay to detect SARS-CoV-2 RNA in FFPE cell pellets using three probes: two probes binding the SARS-CoV-2 positive-sense (genomic) RNA and one probe binding the negative-sense (replicative intermediate) RNA (Figure 2A–F, Supplemental Table 2). As expected, the forty ZZ positive-sense RNA probe 2 binding to SARS-CoV-2 positive-sense RNA resulted in a stronger signal than the twenty ZZ positive-sense RNA probe 1 (Figure 2A– D). Interestingly, in contrast to the wide cytoplasmic distribution of SARS-CoV-2 positive-sense RNA (Figure 2B and D), SARS-CoV-2 negative-sense (replicative intermediate) RNA, detected using negative-sense RNA probe 1 was more specifically localized in perinuclear inclusion bodies (Figure 2F).

**Figure 2.**
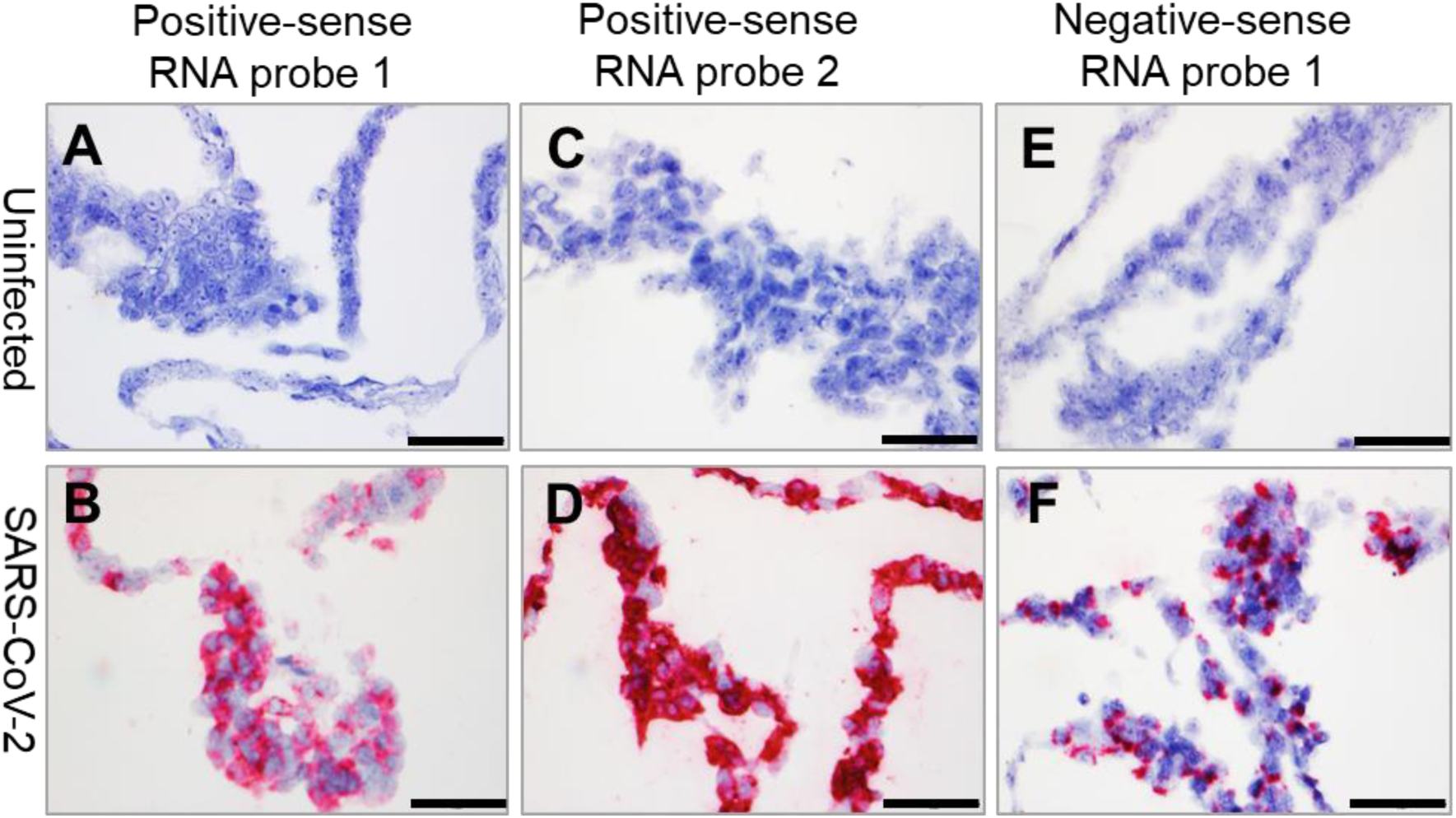
Detection of SARS-CoV-2 RNA by ISH in FFPE cell pellets. (**A–B**) SARS-CoV-2 positive-sense RNA can be detected by ISH using positive-sense RNA probe 1 in infected FFPE cell pellets (**B**), but not in uninfected control FFPE cell pellets (**A**). (**C–D**) SARS-CoV-2 positive-sense RNA can be detected by ISH using positive-sense RNA probe 2 in infected FFPE cell pellets (**D**), but not in uninfected control FFPE cell pellets (**C**). (**E–F**) SARS-CoV-2 negative-sense RNA can be detected by ISH using negative-sense RNA probe 1in infected FFPE cell pellets (**E**), but not in uninfected control FFPE cell pellets (**F**). Nuclei are stained blue (hematoxylin). Scale bar, 50 μm in (**A**–**F**).

### Detection of SARS-CoV-2 replication in FFPE specimens using mFISH

Single-stranded RNA viruses, such as SARS-CoV-2, have to generate a replicative intermediate RNA as a template to synthesize progeny genomic RNAs. We have previously reported the use of mFISH to detect EBOV, MARV, and NiV replication in FFPE tissues (26, 28, 29). Here, we tested mFISH to detect SARS-CoV-2 replication in FFPE specimens using positive-sense RNA probe 2 and negative-sense RNA probe 2 (Supplemental Table 2). Consistent with the RNAscope ISH results, positive-sense viral RNA was widely distributed in the cytoplasm, whereas negative-sense RNA (replicative intermediate) was confined to perinuclear inclusion bodies (Figure 3A– B).

**Figure 3.**
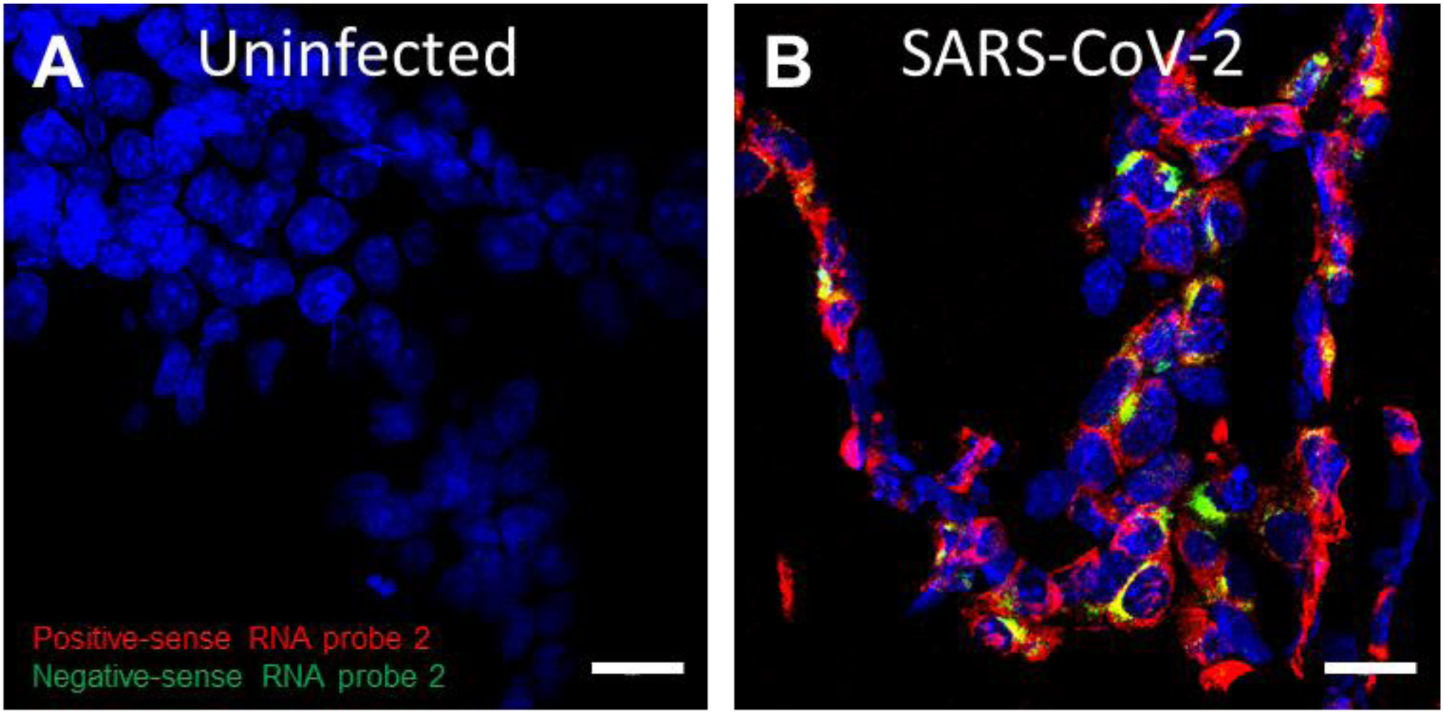
Detection of SARS-CoV-2 replication in FFPE cells using multiplex fluorescence ISH. (**A–B**) Compared to uninfected control (**A**), SARS-CoV-2 negative-sense RNA (green), a replicative intermediate that indicates viral replication, can be detected in infected FFPE cell pellets in addition to positive-sense (red) RNA (**B**). Nuclei are stained blue (DAPI). Scale bar, 20 μm in (**A**–**B**).

### Dual staining to detect SARS-CoV-2 antigen and RNA in the same FFPE section

To more precisely detect SARS-CoV-2, we developed a dual staining assay to recognize both SARS-CoV-2 antigen and RNA in the same FFPE section. IHC was performed using the identified rabbit polyclonal anti-SARS-CoV S antibody following ISH using positive-sense RNA probe 2. Consistently, SARS-CoV-2 antigen was detected along with positive-sense RNA in the cytoplasm of most of the infected, but not in uninfected, cells (Figure 4A–B).

**Figure 4.**
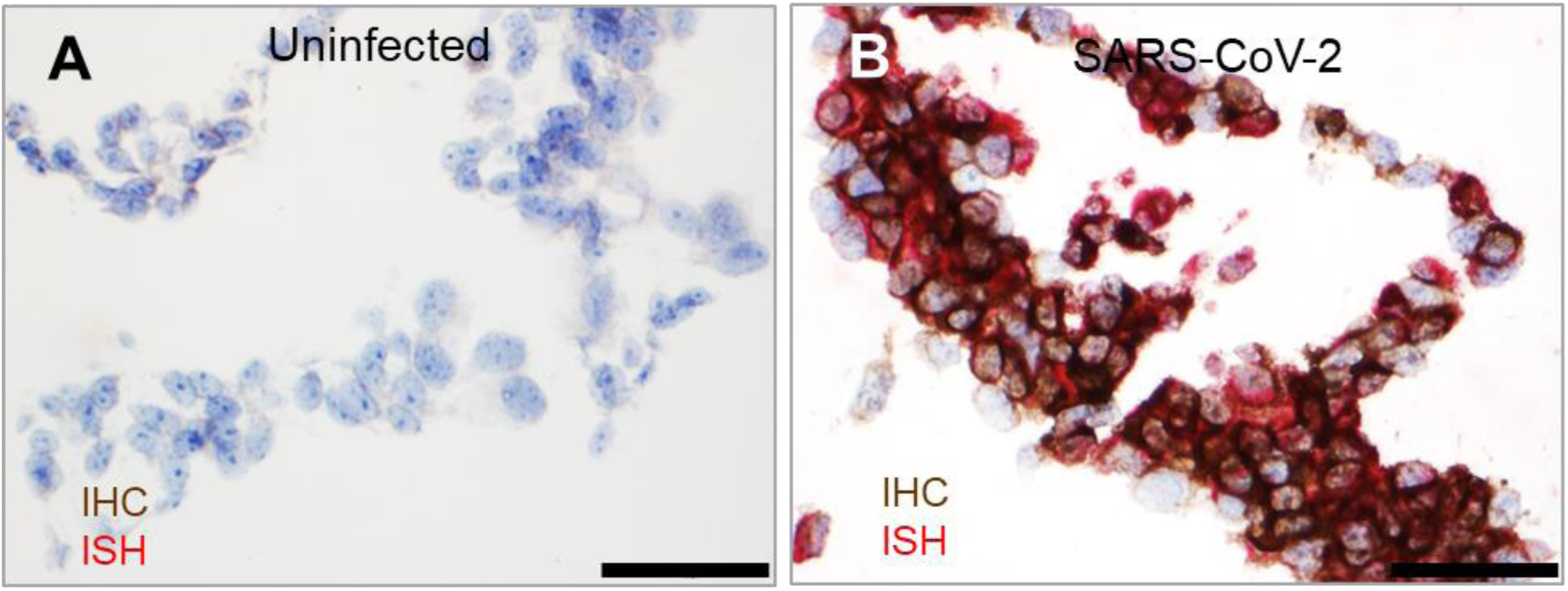
Dual staining to detect SARS-CoV-2 antigen and RNA in the same FFPE section. (**A–B**) Compared to uninfected control FFPE cell pellets (**A**), SARS-CoV-2 S (brown) and positive-sense RNA (red) were detected in the same section (**B**). Nuclei are stained blue (hematoxylin). Scale bar, 50 μm in (**A**–**B**).

## Discussion

As worldwide infectious disease researchers are racing to understand the pathogenesis of and to develop and evaluate MCMs against COVID-19 to contain the ongoing pandemic, assays that determine SARS-CoV-2 distribution in tissues and specific cellular targets of infection are urgently needed. Here we evaluated commercial reagents and assays to detect SARS-CoV-2 antigens or RNA in FFPE specimens. We identified one rabbit polyclonal antibody and one mouse monoclonal antibody that reacts with SARS-CoV-2 S and N, respectively, and demonstrated that these two antibodies can be used to detect SARS-CoV-2 by IHC and IFA in FFPE specimens. Additionally, we characterized two RNAscope ISH assays that can be used to detect SARS-CoV-2 positive- and negative-sense RNAs in FFPE specimens. Furthermore, we developed a dual staining assay using IHC and ISH to detect SARS-CoV-2 S and RNA in the same FFPE section. These reagents and assays are all commercially available and therefore can be applied readily to detect SARS-CoV-2 in both human and animal FFPE tissues.

Detection of viral antigen by IHC and IFA has been widely used to detect infection of high-consequence viruses, including SARS-CoV, EBOV, MARV, LASV, and NiV in human and animal FFPE tissues (26, 28-32). Although various antigen retrieval methods can help to restore the immunoreactivity of epitopes in FFPE tissues, in our experience it remains more challenging to identify antibodies that binds their targets in FFPE tissues compared to frozen section tissues. The FFPE specimen-compatible rabbit and mouse anti-SARS-CoV-2 antibodies we characterized here can be used to map the cellular targets of SARS-CoV-2 in various organs using multiplex IFA in addition to detecting viral infection. RNAscope ISH is a relatively novel ISH platform with high-sensitivity and low-background due to its unique “ZZ” probe design (33). This platform has been widely used to detect viruses both in human and animal tissues (27, 34-36). Single-stranded RNA viruses have to produce a replicative intermediate, antigenomic RNA, as a template to synthesize new genomic RNAs. Presence of such replicative intermediate RNA in tissues indicates ongoing viral replication (26, 28, 29). The commercially available RNAscope ISH assays, including chromogenic and fluorescent assays, we characterized here, can be applied to detect viral RNA in both human and animal tissue samples. The dual staining we developed to detect SARS-CoV-2 viral antigen and RNA in the same FFPE section can more precisely detect SARS-CoV-2 because a positive IHC or ISH signal alone may originate from remaining free viral antigen or degenerating RNA fragments rather than from viral particles. Because SARS-CoV-2-infected animal tissues were not available at the time of this study, we were restricted to evaluate FFPE pellets of Vero 76 cells as a surrogate. However, we prepared FFPE cell pellets using the same process used for FFPE tissues preparation. Additionally, FFPE cell pellets have been widely used to evaluate antibodies and ISH assays and other reagents for FFPE tissue analysis and have been largely predictive of reactivity with genuine tissues (33, 37, 38). We are therefore confident that the SARS-CoV-2 IHC, ISH, mFISH, and dual staining assays we developed and characterized will be highly useful to study pathogenesis of SARS-CoV-2 infection in both human and animal models.

## Methods

### Cells and virus

Grivet (*Chlorocebus aethiops*) Vero 76 kidney epithelial cells (American Type Culture Collection [ATCC], Manassas, VA; #CRL-1587) were maintained in Eagle’s minimum essential media (EMEM; Thermo Fisher Scientific, Waltham, MA, USA) supplemented with 10% heat-inactivated FBS (Hyclone, Logan, UT, USA), 1% GlutaMax (Thermo Fisher Scientific) and1% non-essential amino acid solution (MilliporeSigma, Temecula, CA, USA), at 37°C in a 5% CO2 atmosphere. The SARS-CoV-2 USA-WA1/2020 strain (GenBank #MN985325.1) was obtained from the US Centers for Disease Control and Prevention (CDC, Atlanta, GA, USA). Virus was added to Vero 76 cell cultures in T-75 flasks in biosafety level 3 (BSL-3) containment at a multiplicity of infection (MOI) of 0.01. Cells were then incubated for 1 h for virus adsorption, washed with EMEM, and maintained in EMEM with 10% FBS. Cells were fixed at 24 h post-inoculation in 10% neutral buffered formalin for 24 h and then moved from the BSL-3 to a BSL-2 suite. Uninfected Vero 76 were processed as a control.

### Cell pellet embedding

Fixed cells were scraped off flasks after being rinsed twice in PBS (Thermo Fisher Scientific). Scraped cells were spun down at 2,500 rpm and the pellets were mixed with liquefied Histogel (Thermo Fisher Scientific). Pellets were solidified at 4°C and further processed for paraffin embedding using an automated Tissue Tek VIP processor (Sakura Finetek, Torrance, CA, USA).

### IHC

IHC was performed using the Envision system (Dako Agilent Pathology Solutions, Carpinteria, CA, USA). Briefly, after deparaffinization, peroxidase blocking, and antigen retrieval, sections were covered with a primary antibody at a 1;1000, 1:2000, or 1:4000 dilution (Supplemental Table 1) and incubated at room temperature for 45 min. Subsequently, sections were rinsed, and the peroxidase-labeled polymer (secondary antibody) was applied for 30 min. Slides were rinsed and a brown chromogenic substrate DAB solution (Dako Agilent Pathology Solutions) was applied for 8 min. The substrate-chromogen solution was rinsed off the slides, and slides were counterstained with hematoxylin and rinsed. The sections were dehydrated, cleared with Xyless II (Valtech, Brackenridge, PA, USA), and then coverslipped.

### IFA

After deparaffinization and reduction of autofluorescence, tissues were heated in citrate buffer, pH 6.0 (MilliporeSigma), for 15 min to reverse formaldehyde cross-links. After rinsing with PBS, pH 7.4 (Thermo Fisher Scientific), sections were blocked overnight with CAS-Block (Thermo Fisher Scientific) containing 5% normal goat serum (MilliporeSigma) at 4°C. Sections were then incubated with rabbit polyclonal antibody against SARS-CoV S (Sino Biological, #40150-T62-COV2) at dilution 1:500 and mouse monoclonal antibody against SARS-CoV NP (Sino Biological, 40143-MM05) at a dilution 1:500 for 2 h at room temperature. After rinsing in PBST (PBS + 0.1% Tween-20, MilliporeSigma), sections were incubated with secondary goat IgG Alexa Fluor 488-conjugated anti-rabbit antibody and with goat IgG Alexa Fluor 561-conjugated anti-mouse antibody (Thermo Fisher Scientific) for 1 h at room temperature. Sections were cover-slipped using VECTASHIELD antifade mounting medium with DAPI (Vector Laboratories, Burlingame, CA, USA). Images were captured on an LSM 880 Confocal Microscope (Zeiss, Oberkochen, Germany) and processed using open-source ImageJ software (National Institutes of Health, Bethesda, MD, USA).

### ISH

To detect SARS-CoV-2 genomic RNA in FFPE tissues, ISH was performed using the RNAscope 2.5 HD RED kit (Advanced Cell Diagnostics, Newark, CA, USA) according to the manufacturer’s instructions. Briefly, forty ZZ ISH probes (#854841, positive-sense RNA probe 1) with C1 channel and twenty ZZ ISH probes (#848561, positive-sense RNA probe 2) with C1 channel targeting SARS-CoV-2 positive-sense (genomic) RNA and twenty ZZ ISH probes (#845701, negative-sense RNA probe 1) with C1 channel targeting SARS-CoV-2 negative-sense (replicative intermediate) RNA were designed and synthesized by Advanced Cell Diagnostics (Supplemental Table 2). Tissue sections were deparaffinized with xylene, underwent a series of ethanol washes and peroxidase blocking, and were then heated in kit-provided antigen retrieval buffer and digested by kit-provided proteinase. Sections were exposed to ISH target probe pairs and incubated at 40°C in a hybridization oven for 2 h. After rinsing, ISH signal was amplified using kit-provided Pre-amplifier and Amplifier conjugated to alkaline phosphatase and incubated with a Fast Red substrate solution for 10 min at room temperature. Sections were then stained with hematoxylin, air-dried, mounted, and stored at 4°C until image analysis.

### Multiplex fluorescence ISH

Multiplex fluorescence ISH (mFISH) was performed using the RNAscope Fluorescent Multiplex Kit (Advanced Cell Diagnostics) according to the manufacturer’s instructions with minor modifications. In addition to positive-sense RNA probe 1 (red), another forty ZZ probes with C3 Channel (green, #854851-C3, negative-sense RNA probe 2) targeting negative-sense (replicative intermediate) SARS-CoV-2 RNA was designed and synthesized by Advanced Cell Diagnostics (Supplemental Table 2). FFPE-tissues sections underwent deparaffinization with xylene and a series of ethanol washes and treatment with 0.1% Sudan Black B (Sigma-Aldrich, St. Louis, MO, USA) to reduce autofluorescence. Tissues were heated in kit-provided antigen retrieval buffer and digested by kit-provided proteinase. Sections were exposed to mFISH target probes and incubated at 40°C in a hybridization oven for 2 h. After rinsing, mFISH signal was amplified using company-provided Pre-amplifier and Amplifier conjugated to fluorescent dye. Sections were counterstained with DAPI (Thermo Fisher Scientific), mounted, and stored at 4°C until image analysis. mFISH images were captured on an LSM 880 Confocal Microscope (Zeiss, Oberkochen, Germany) and processed using open-source ImageJ software (National Institutes of Health, Bethesda, MD, USA).

### Dual staining

Sections were covered with rabbit polyclonal anti-SARS-CoV S antibody diluted at 1:250 (Sino Biologicals, #40150-T62-COV2, Supplemental Table 1) overnight at 4°C, following the Fast Red substrate ISH procedure described above using positive-sense RNA probe 2 (Supplemental Table 2). One day later, sections were rinsed, and the peroxidase-labeled polymer (secondary antibody) was applied for 45 min. Slides were rinsed and a brown chromogenic substrate, 3,3’-iaminobenzidine (DAB) solution (Dako Agilent Pathology Solutions), was applied for 8 min. Sections were then stained with hematoxylin, air-dried, and mounted, and stored at 4°C until image analysis.

## Author contributions

X.Z. conceived and designed the experiments. J. L., A.M.B., B.J.K., S.R.R., and X.Z. performed experiments. S.R.R., J. H.K., and X.Z. interpreted the data and wrote the manuscript with input from all authors.

## Acknowledgments

We thank Lynda Miller and Neil Davis (USAMRIID, Frederick, MD, USA) for histology assistance and Paul Facemire and Kathleen Gibson at USAMRIID for coordinating the cell infection study.

This work was funded by Defense Health Program (DHP). JHK’s participation was funded, in part, through Laulima Government Solutions, LLC prime contract with the US National Institute of Allergy and Infectious Diseases (NIAID) under Contract No. HHSN272201800013C. J.H.K. performed this work as an employee of Tunnell Government Services (TGS), a subcontractor of Laulima Government Solutions, LLC under Contract No. HHSN272201800013C. The views and conclusions contained in this document are those of the authors and should not be interpreted as necessarily representing the official policies, either expressed or implied, of the US Department of the Army, the US Department of Defense, the US Department of Health and Human Services, or of the institutions and companies affiliated with the authors. In no event shall any of these entities have any responsibility or liability for any use, misuse, inability to use, or reliance upon the information contained herein. The US departments do not endorse any products or commercial services mentioned in this publication..

## Conflict of interest statement

The authors have declared that no conflict of interest exists.

## Supplemental Tables

**Supplemental Table 1.** Antibodies tested for detecting SARS-CoV-2 in FFPE specimens.

| **Antibody** | **Source** | **Catalog**<br>**number** | **Immunogen** | **Clonality** | **IHC Suitability**<br>**(FFPE**<br>**Specimen)** |
| --- | --- | --- | --- | --- | --- |
| Rabbit anti-SARS-CoV<br>spike<br>glycoprotein<br>(S) | Novus<br>Biologicals,<br>Littleton, CO,<br>USA | NB100-56048 | SARS-CoV S residues<br>288–303 | Polyclonal | No |
| Rabbit anti-SARS-CoV<br>spike<br>glycoprotein<br>(S) | Novus<br>Biologicals,<br>Littleton, CO,<br>USA | NB100-56578 | SARS-CoV S residues<br>1124–1140 | Polyclonal | No |
| Rabbit anti-SARS-CoV-2 spike glycoprotein (S) | Sino Biological, Chesterbrook, PA, USA | 40150-R007 | Recombinant SARS-CoV-2 S subunit S1 | Monoclonal | No |
| Rabbit anti-SARS-CoV spike glycoprotein (S) | Sino Biological, Chesterbrook, PA, USA | 40150-T62-COV2 | Recombinant SARS-CoV S subunit S1 | Polyclonal | Yes |
| Rabbit anti-SARS-CoV nucleocapsid protein (NP) | Sino Biological, Chesterbrook, PA, USA | 40143-R001 | Recombinant SARS-CoV NP | Monoclonal | No |
| Rabbit anti-SARS-CoV nucleocapsid protein (NP) | Thermo Fisher Scientific, Waltham, MA, USA | PA1-41098 | SARS-CoV NP residues 399–411 | Polyclonal | No |
| Mouse anti-SARS-CoV nucleocapsid protein (NP) | Sino Biological, Chesterbrook, PA, USA | 40143-MM05 | Recombinant SARS-CoV NP | Monoclonal | Yes |

**Supplemental Table 2.**
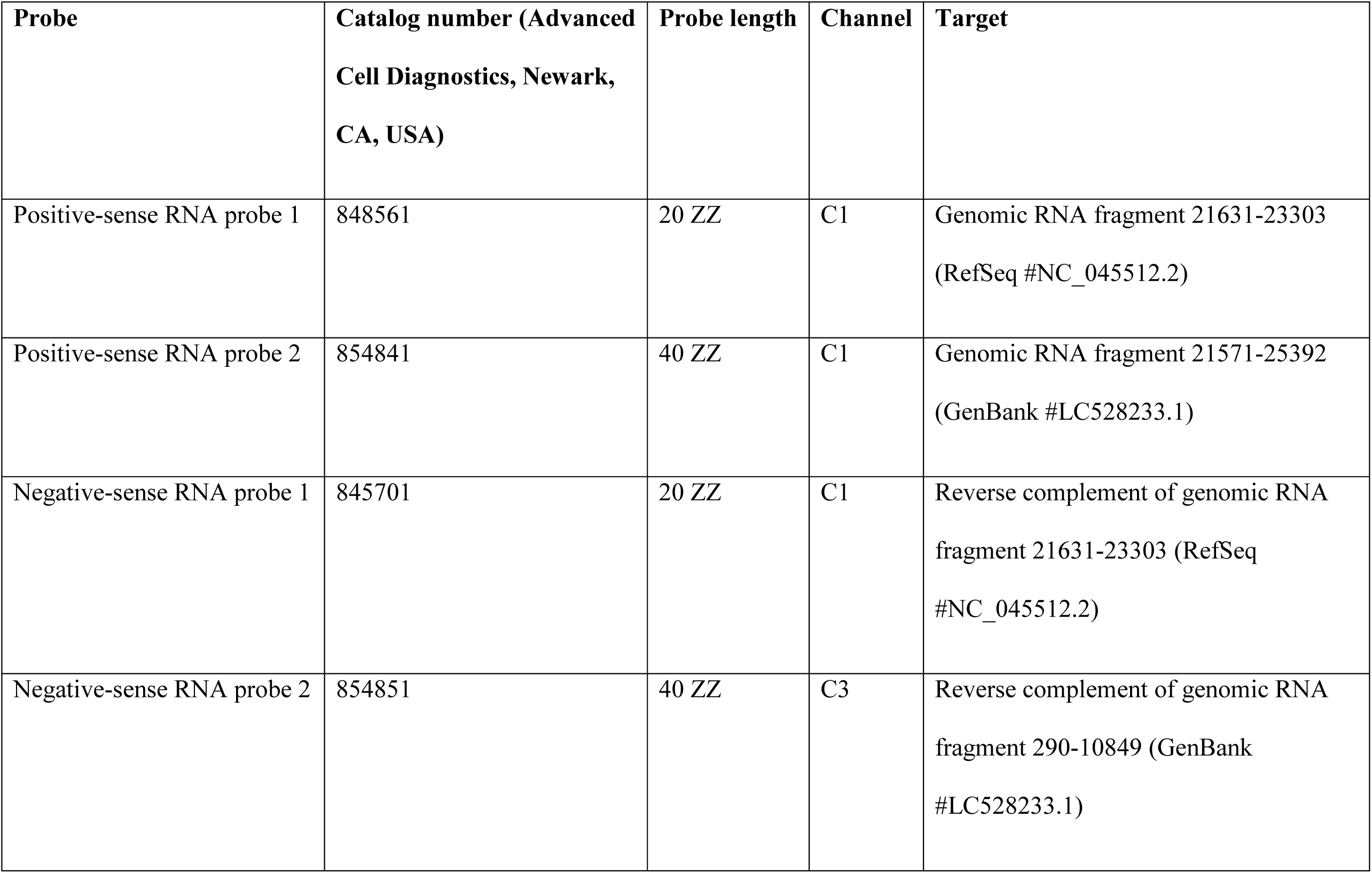
RNAscope ISH probes suitable for detecting SARS-CoV-2 RNA in FFPE specimens.

## Notes

### Competing Interest Statement

The authors have declared no competing interest.

